# Targeting the menopause transition with metformin improves breast cancer outcomes, but discontinuation has deleterious effects on metabolic health: Findings from a preclinical model of postmenopausal breast cancer

**DOI:** 10.1101/2024.11.25.625083

**Authors:** Karen A. Corleto, Pepper Schedin, Anuhya S. Kotta, Jenna L. Strandmo, Samantha M. Foster, Nanci Lammoglia, Moumita Karmakar, Raymond J. Carroll, Paul S. MacLean, Erin D. Giles

## Abstract

**Background:** Women with obesity and/or type-II-diabetes have an increased breast cancer risk, increased metastasis, and poorer prognosis, especially after menopause. In a rat model of high-fat-diet and menopause-induced weight gain, we previously reported that treatment with the anti-diabetic drug metformin for 8-weeks after ovariectomy (OVX; modeling menopause) reduced growth of existing mammary tumors and inhibited new tumor formation. This identified the menopause transition as a potential window-of-opportunity for interventions to decrease obesity-associated breast cancer incidence and disease progression. Here, we extend these findings to determine if limiting metformin to the peak window of OVX-induced weight gain would have similar anti-cancer effects.

**Findings:** Metformin during the first four weeks following OVX is critical to reducing tumor burden, as rats treated with metformin early (weeks0-4-postOVX) had reduced tumor burden. Conversely, initiating metformin later in the postOVX period (weeks 4-8postOVX) did not reduce cancer burden. Despite improved tumor outcomes, metformin withdrawal after the early postOVX time had detrimental metabolic effects, including weight gain and increased adiposity, insulin, IGF1, and HOMA-IR, which correlate with increased cancer risk.

**Conclusions:** These data reveal early-postmenopause as a critical window when metformin decreases progression of existing disease and highlights the importance of maintaining treatment to prevent metabolic dysregulation, which could promote secondary tumors/metastasis. These findings also help explain the disconnect between epidemiological studies reporting anticancer benefits of metformin and more recent clinical trials that failed to see similar efficacy, potentially due to issues of timing and/or inclusion of women outside the early postmenopausal window and/or without underlying metabolic dysfunction.

## BACKGROUND

Obesity and type-II-diabetes (T2D) increase breast cancer (BC) risk and poor clinical outcomes[1, 2]. During menopause, obesity-associated pathophysiological changes occur, including weight gain, hormonal dysregulation, increased central adiposity, and chronic inflammation, which can exacerbate the tumor-promoting effects of obesity[2, 3]. Metformin, a conventional anti-diabetic drug, decreases BC incidence and mortality in individuals with diabetes when compared to those on other diabetic drugs[4-6], through systemic and tumor-specific effects [6, 7]. Epidemiological data indicate a dose-response relationship where higher doses and longer durations of metformin treatment may be most beneficial[4, 5]. Clinical trials, however, have not shown consistent benefits[8, 9]. Conflicting results may be due to differences in patient metabolic status (diabetes/no diabetes) and/or the timing of treatment. Specifically, the menopause transition may represent a targetable window of opportunity for interventions to decrease obesity-associated cancer risk.

In a rat model of diet-induced obesity and postmenopausal breast cancer, we previously demonstrated that weight gain, increased visceral adiposity, and associated insulin resistance induced by surgical ovariectomy (OVX; model of menopause) increase mammary tumor burden[3, 10-13], and metformin treatment targeted up to 8 weeks post-OVX decreased tumor burden[14]. Our goal in the current study was to build on these findings and determine if limiting metformin to the narrower window of maximum OVX-induced weight gain (i.e., 3-4 weeks post-OVX in rats) is sufficient to recapitulate these anti-tumor effects of metformin.

## METHODS

### Animals

All procedures were IACUC-approved and were identical to our long-term metformin study[14], with two additional treatment groups evaluating the effect of metformin timing (**Fig-1A; Supplemental-Table-1**). Briefly, rats were fed high-fat diet (46%kcal fat) to induce obesity, given a single injection of carcinogen (N-methyl-nitrosourea; MNU) to induce mammary tumors, and surgically ovariectomized (OVX) to model menopause. Rats were randomized to receive metformin during either the early (weeks0-4 postOVX; MET_0-4wks_), late (weeks4-8 postOVX, MET_4-8wks_), or full 8-week postOVX period (MET_0-8wks_). Untreated rats served as controls. Tumor development/growth, body weight, and body composition were measured throughout the study.

**Figure 1.**
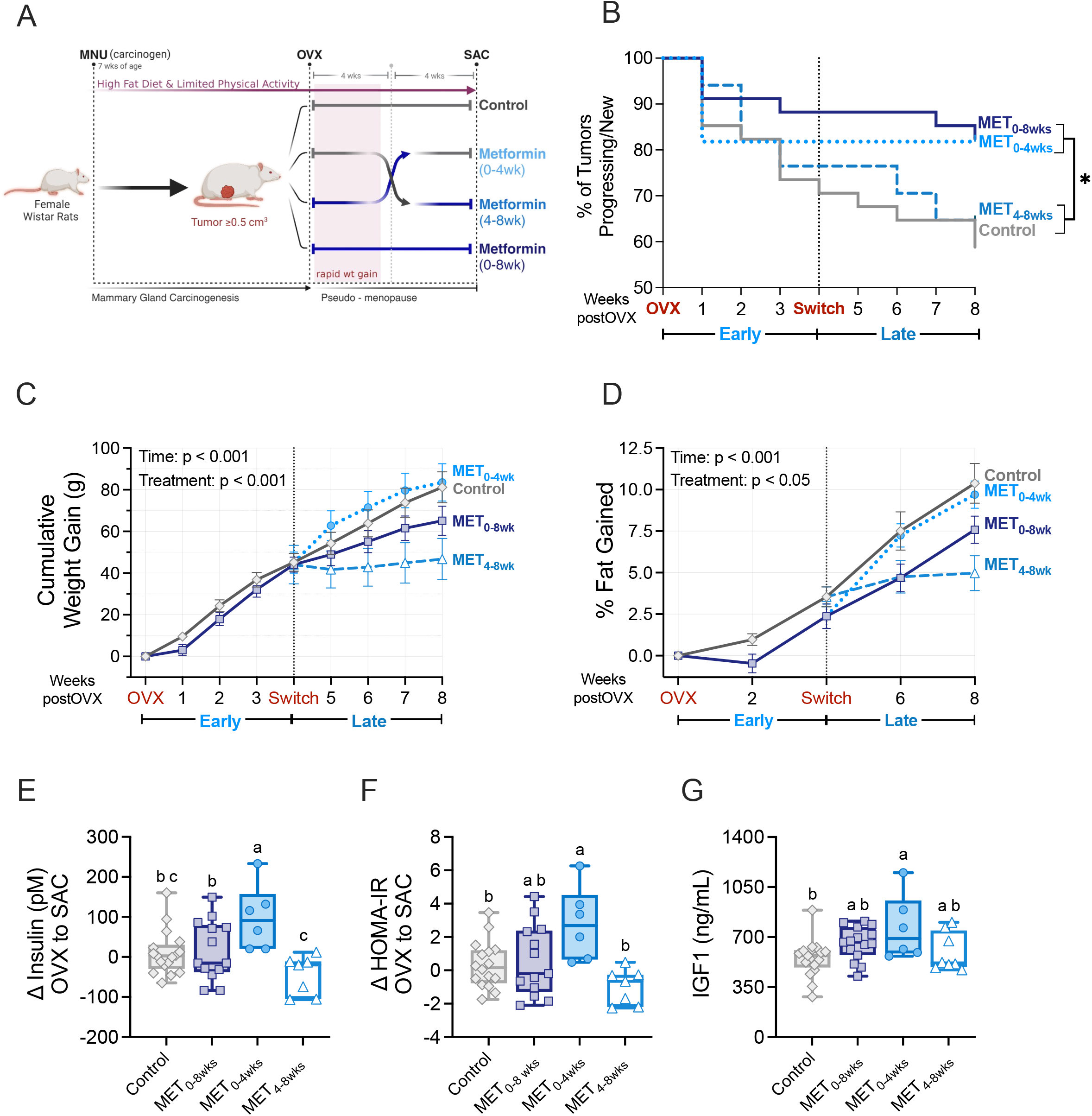
Study design, tumor outcomes, metabolic outcomes, and body composition throughout the study. **A)** Study design and timeline. **B**) Kaplan Meier curve of tumor progression for eight weeks post-ovariectomy (OVX). **C**) Weight gain post-ovariectomy (OVX), as a percent of pre-OVX body weight. **D**) Percent body fat increase biweekly from OVX to study end. Change in **E)** plasma insulin and **F)** HOMA-IR from OVX to study end. **G**) Plasma IGF1 levels at the end of the study. Box and whisker plots show median, minimum, and maximum values. For line graphs, data are represented as mean ± SE. *p < 0.05. ^a,b,c^Groups with the same letter designation are not significantly different (*p* < 0.05).

### Plasma

Tail vein blood was collected at OVX and euthanasia, plasma isolated, and stored at −80°C. Plasma metabolites were measured by ELISA and colorimetric assays (**Supplemental-Table-2)**.

### Histology

To assess lipid accumulation, formalin-fixed, paraffin-embedded (FFPE) livers were stained with adipophilin antibody (LS-C348703, Lifespan-Biosciences, 1:300) as described[12]. Macrophages forming crown-like structures (CLS) were identified by staining FFPE tumors and mammary adipose for CD68 (Ab4059, Serotec, 1:200)[12]. Number of CLS was counted by three independent researchers and averaged (inter-observer SD=3.1 CLS/cm^2^). For adipocyte size, sections of non-tumor-bearing mammary adipose and tumors with adjacent adipose were stained with H&E, scanned at 20X (Aperio slide scanner, Leica-Biosystems), and adipocyte diameter measured in three randomly selected regions of each slide (Adiposoft-plug-in for ImageJ[15]).

### Statistics

Statistical analyses were conducted using R (v4.0.4) and GraphPad Prism (v10). Data were analyzed using T-tests or ANOVA and equivalent non-parametric tests when applicable. Data are presented as mean±standard error. One rat randomized to MET_0-4wks_ was excluded from all analyses due to the development of several off-target tumors.

## RESULTS

### Metformin during, but not after, OVX-induced weight gain decreases tumor progression

Metformin during the early 4-week postOVX window of rapid weight gain reduced tumor progression similar to full 8-weeks of treatment (**Fig-1B)**. In contrast, treating rats from weeks 4-8 postOVX, after the window of rapid OVX-induced weight gain, did not confer protection, suggesting that the timing of metformin treatment relative to loss of ovarian function and associated weight gain is critical in mediating its anti-cancer effects.

### Terminating metformin treatment promotes weight gain and negatively impacts metabolic health

OVX promoted weight gain and increased body fat in all rats during the early (weeks 0-4) postOVX period (**Fig-1C,D**), mirroring weight gain and increased adiposity that are common during menopause and consistent with our previous studies[10, 11, 14], with no difference observed between metformin-treated and control rats. Withdrawal of metformin in the MET_0-4wks_ group increased the trajectory of bodyweight and adipose deposition (effect of treatment and time, p<0.001 for both), while treatment with metformin across weeks 4-8 postOVX (MET_4-8wks_) blunted weight gain and adipose deposition (**Figs-1C-D; Supplemental-Table3**). Similarly, stopping metformin at four weeks postOVX (MET_0-4wks_) resulted in significantly higher circulating insulin, insulin resistance (HOMA-IR), and IGF1 at 8-weeks postOVX (**Fig-1E-G)**. Glucose, cholesterol, triglycerides, non-esterified fatty acids, and leptin did not differ between groups (**Supplemental-Table-4)**.

### Metformin reduces size of tumor-border adipocytes

Given that adipocyte size and inflammation are known to influence insulin sensitivity, we assessed the impact of metformin on these markers of adipose health in nontumor-bearing mammary glands and mammary adipose tissue in the tumor border to evaluate potential differences in the adipose environment depending on its proximity to tumor tissue. In non-tumor mammary tissue, treatment did not impact adipocyte diameter (**Fig-2A,B)**. However, in the tumor border (**Fig-2C**), metformin-treated rats had a higher proportion of very small adipocytes (20-50μm) vs control rats, regardless of timing/duration of metformin. Interestingly, in the MET_0-4wks_ rats, the proportion of large adipocytes (>90um) was similar to control rats, which parallels their increased adiposity after metformin withdrawal. Together, these differences resulted in significantly lower mean adipocyte diameter in the MET_0-8wks_ and MET_4-8wks_ groups (**Fig-2D)**. A hallmark of adipose inflammation is the presence of crown-like structures (CLS), formed when macrophages engulf dying adipocytes. Paralleling the decreased adipocyte size, continuous (MET_0-8wks_) and late (MET_4-8wks_) metformin treatment tended to reduce the number of CLS in the tumor border compared to controls (p<0.1; **Fig-2E,F**).

**Figure 2.**
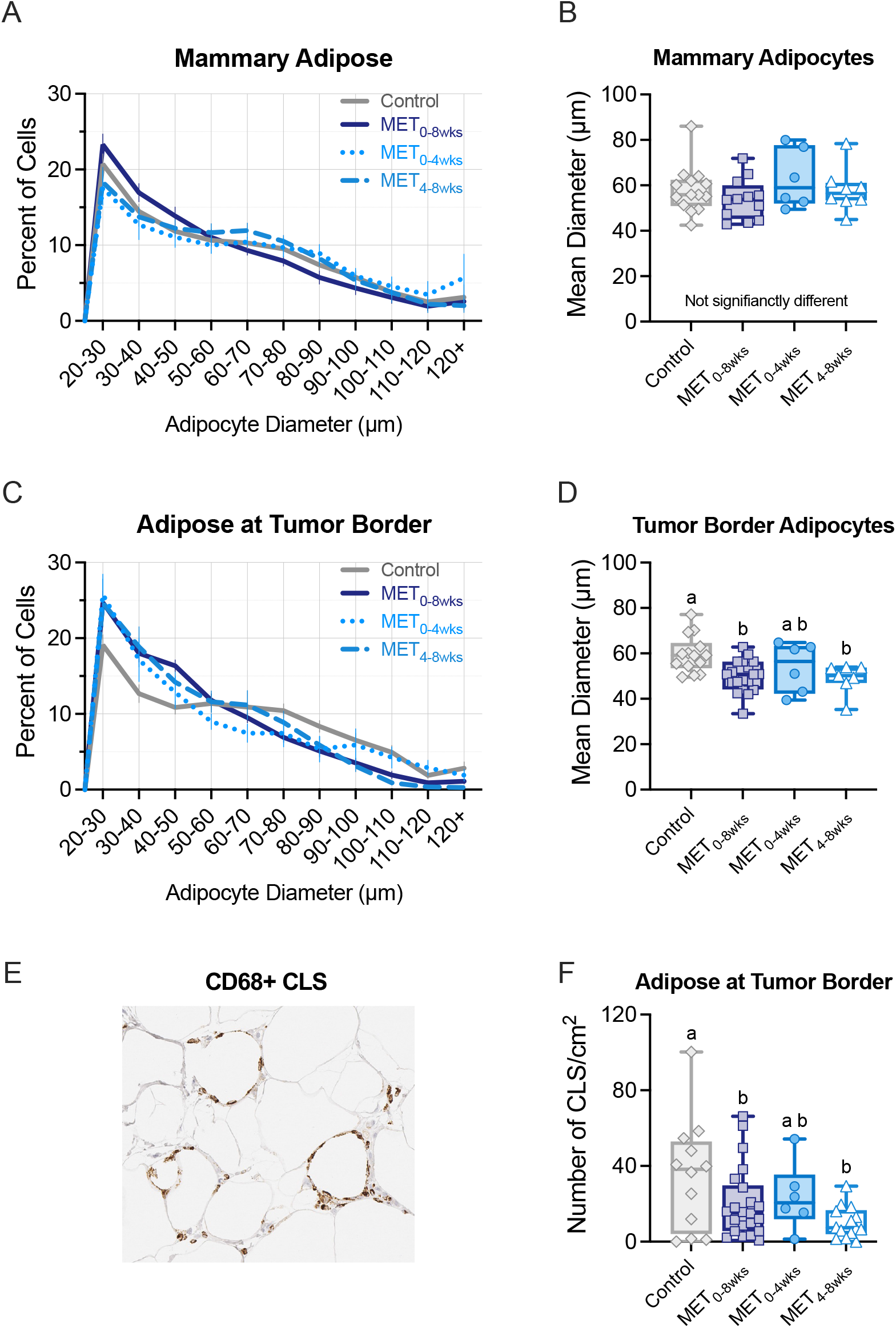
The effect of metformin timing on adipocyte size and inflammation. Size distribution and **B)** mean diameter of adipocytes from subcutaneous/mammary gland of control, MET_0-8wks_, MET_0-4wks_, and MET_4-8wks_ rats, measured at study end. **C)** Size distribution and **D)** mean diameter of adipocytes from adipose surrounding the mammary tumor border of control, MET_0-8wks_, MET_0-4wks_, and MET_4-8wks_ rats, measured at study end. **E)** Representative image of IHC staining for CD68+ indicating the presence of crown-like structures (CLS) in the mammary tumor border (10x magnification). **F)** Number of CLS/cm^2^ in the mammary tumor border. *p < 0.05; ^a,b^Groups with the same letter designation are not significantly different (*p* < 0.05).

## DISCUSSION

Menopause is a critical lifecycle window of BC risk, where changes in energy balance, hormones, chemokines/cytokines, and increased adiposity are pro-tumorigenic. We previously reported that long-term metformin treatment, initiated at the time of OVX (to model menopause), reduced tumor burden, in part due to a reduction in aromatase-positive macrophages in the tumor microenvironment[14]. Despite this and other preclinical studies showing beneficial effects, findings from clinical intervention studies of metformin remain ambiguous, with the recent MA.32 trial failing to see improvement in cancer risk or overall survival[8, 9]. In support of our previous hypothesis[6], our current study demonstrates that both the timing and duration of metformin treatment, particularly relative to ovariectomy, are critical to its anti-cancer potential. These data highlight the early postmenopausal window as the time when metformin is likely to have the greatest anti-cancer activity, whereas initiating treatment later in the postmenopausal period could be insufficient to reduce tumor incidence or progression.

These data also suggest that metabolic impairment, such as that induced by OVX and HFD, may be necessary for metformin’s anti-cancer activity. MA.32, a key clinical study assessing metformin’s anti-cancer potential, was limited to women without type-II-diabetes or metabolic dysfunction[8, 9], potentially explaining the lack of efficacy in this trial. Preclinically, studies using diets high in fat and/or sucrose to induce metabolic disease have reported anti-cancer benefits with metformin[14, 16-18], whereas those using low-fat diets have reported negligible impact on tumors[19, 20]. While diet composition was not evaluated in any of the clinical cancer studies assessing metformin, metabolic syndrome has been associated with “unhealthy” dietary patterns[21], suggesting that those without metabolic disease may be more likely to consume a “healthy” diet and thus not see the same benefits from metformin treatment.

Our data also indicate that targeting the menopausal window, a time highly associated with metabolic dysfunction, may be key to optimizing the efficacy of metformin as a breast cancer preventive agent. Corroborative human data shows that most of the metformin-mediated reductions in BC mortality have been seen in early-stage BC[22], suggesting that metformin may be most effective in the prevention setting.

Another key finding from our study is that abrupt discontinuation of metformin was detrimental to metabolic health, with a significant rise in insulin, IGF1, and markers of insulin resistance all linked to increased risk and reduced survival in patients with BC[23]. While few studies have examined the discontinuation of metformin in a cancer setting, a case-control study in patients with BC and history of metformin use supports this finding, reporting increased risk of both overall and BC-specific mortality within the first 12 months of discontinuing metformin[24].

We find that both continuous and late metformin treatment reduced adipocyte size and tended to lower the number of inflammatory crown-like structures, adipocyte attributes expected to reduce cancer risk[25]. Conversely, adipocyte size increased in the early metformin group and CLS number appeared to be increasing upon metformin discontinuation, further highlighting the potential negative consequences of stopping metformin treatment. These effects were limited to tumor-adjacent adipose, suggesting tumor-specific interactions between metformin and tumor-secreted factors that can modulate the tumor microenvironment. Metformin’s effects on the tumor microenvironment were similarly observed in our previous study where long-term metformin decreased aromatase-expressing macrophages specifically in the tumor border[14].

## CONCLUSION

In summary, the current results demonstrate that metformin treatment during the first four weeks post-OVX effectively decreases tumor progression, yet stopping metformin early leads to negative metabolic outcomes and increased cancer risk. Our findings also indicate that when metformin is initiated after the peak window of OVX-induced weight gain, it does not effectively mitigate tumor progression. Together, this suggests that the timing of metformin treatment during the menopause transition is critical for tumor responses to treatment, with increased efficacy when initiated in perimenopause or early menopause, and maintaining treatment throughout this lifecycle window is likely necessary for maintenance of metabolic health.

## LIST OF ABBREVIATIONS

AT: adipose tissue
BC: breast cancer
CLS: crown-like structures
ER+: estrogen receptor-positive
FIJI: ImageJ
H&E: hematoxylin and eosin
HOMA-IR: homeostatic model assessment for insulin resistance
IGF-1: insulin-like growth factor 1
IHC: immunohistochemistry
MET: metformin
MET_0-8wks_: metformin treatment for eight weeks
MET_0-4wks_: metformin treatment for the first four weeks of the eight-week period
MET_4-8wks_: metformin treatment for the last four weeks of an eight-week period
MG: mammary gland
MNU: 1-methyl-1-nitrosourea
NEFA: non-esterified fatty acids – plasma-free fatty acids
OVX: ovariectomized
qMR: quantitative magnetic resonance
SEM: standard error of the mean
TG: triglycerides

## DECLARATIONS

### Ethics approval

All animal procedures were approved by the University of Colorado Anschutz Medical Campus and the Texas A&M University Institutional Animal Care and Use Committees.

### Consent for publication

Not applicable

## Availability of data and materials

The datasets used and/or analyzed during the current study are available from the corresponding author upon reasonable request.

## Authors’ contributions

KAC participated in data analysis and interpretation. ASK, JLS, SMF, and NL analyzed histological samples. MK and RJC assisted in the statistical analysis and interpretation of the data. PS, PSM, and EDG oversaw all aspects of study design, execution, and data analysis/interpretation. KAC and EDG wrote the manuscript. All authors read and approved the final manuscript.

## Acknowledgments

The study design figure was created using BioRender.com.

## REFERENCES

1. Sun X, Zhang Q, Kadier K, Hu P, Liu X, Liu J, Yan Y, Sun C, Yau V, Lowe S et al: Association between diabetes status and breast cancer in US adults: findings from the US National Health and Nutrition Examination Survey. Front Endocrinol (Lausanne) 2023, 14:1059303.

2. Kim DS, Scherer PE: Obesity, Diabetes, and Increased Cancer Progression. Diabetes Metab J 2021, 45(6):799–812.

3. Martinson HA, Lyons TR, Giles ED, Borges VF, Schedin P: Developmental windows of breast cancer risk provide opportunities for targeted chemoprevention. Experimental cell research 2013, 319(11):1671–1678.

4. Evans JM, Donnelly LA, Emslie-Smith AM, Alessi DR, Morris AD: Metformin and reduced risk of cancer in diabetic patients. Bmj 2005, 330(7503):1304–1305.

5. Bodmer M, Meier C, Krähenbühl S, Jick SS, Meier CR: Long-term metformin use is associated with decreased risk of breast cancer. Diabetes care 2010, 33(6):1304–1308.

6. Corleto KA, Strandmo JL, Giles ED: Metformin and Breast Cancer: Current Findings and Future Perspectives from Preclinical and Clinical Studies. Pharmaceuticals (Basel) 2024, 17(3).

7. Heckman-Stoddard BM, DeCensi A, Sahasrabuddhe VV, Ford LG: Repurposing metformin for the prevention of cancer and cancer recurrence. Diabetologia 2017, 60(9):1639–1647.

8. Goodwin PJ, Chen BE, Gelmon KA, Whelan TJ, Ennis M, Lemieux J, Ligibel JA, Hershman DL, Mayer IA, Hobday TJ et al: Effect of Metformin Versus Placebo on New Primary Cancers in Canadian Cancer Trials Group MA.32: A Secondary Analysis of a Phase III Randomized Double-Blind Trial in Early Breast Cancer. J Clin Oncol 2023:Jco2300296.

9. Goodwin PJ, Chen BE, Gelmon KA, Whelan TJ, Ennis M, Lemieux J, Ligibel JA, Hershman DL, Mayer IA, Hobday TJ et al: Effect of Metformin vs Placebo on Invasive Disease-Free Survival in Patients With Breast Cancer: The MA.32 Randomized Clinical Trial. Jama 2022, 327(20):1963–1973.

10. Giles ED, Wellberg EA: Preclinical Models to Study Obesity and Breast Cancer in Females: Considerations, Caveats, and Tools. Journal of Mammary Gland Biology and Neoplasia 2020, 25(4):237–253.

11. Giles ED, Jackman MR, Johnson GC, Schedin PJ, Houser JL, MacLean PS: Effect of the estrous cycle and surgical ovariectomy on energy balance, fuel utilization, and physical activity in lean and obese female rats. Am J Physiol Regul Integr Comp Physiol 2010, 299(6):R1634–1642.

12. Wellberg EA, Corleto KA, Checkley LA, Jindal S, Johnson G, Higgins JA, Obeid S, Anderson SM, Thor AD, Schedin PJ et al: Preventing ovariectomy-induced weight gain decreases tumor burden in rodent models of obesity and postmenopausal breast cancer. Breast Cancer Research 2022, 24(1):42.

13. Sherk VD, Jackman MR, Giles ED, Higgins JA, Foright RM, Presby DM, Johnson GC, Houck JA, Houser JL, Oljira R et al: Prior weight loss exacerbates the biological drive to gain weight after the loss of ovarian function. Physiol Rep 2017, 5(10):e13272.

14. Giles ED, Jindal S, Wellberg EA, Schedin T, Anderson SM, Thor AD, Edwards DP, MacLean PS, Schedin P: Metformin inhibits stromal aromatase expression and tumor progression in a rodent model of postmenopausal breast cancer. Breast Cancer Res 2018, 20(1):50.

15. Galarraga M, Campión J, Muñoz-Barrutia A, Boqué N, Moreno H, Martínez JA, Milagro F, Ortiz-de-Solórzano C: Adiposoft: automated software for the analysis of white adipose tissue cellularity in histological sections. J Lipid Res 2012, 53(12):2791–2796.

16. Checkley LA, Rudolph MC, Wellberg EA, Giles ED, Wahdan-Alaswad RS, Houck JA, Edgerton SM, Thor AD, Schedin P, Anderson SM: Metformin accumulation correlates with organic cation transporter 2 protein expression and predicts mammary tumor regression in vivo. Cancer Prevention Research 2017, 10(3):198–207.

17. Orecchioni S, Reggiani F, Talarico G, Mancuso P, Calleri A, Gregato G, Labanca V, Noonan DM, Dallaglio K, Albini A et al: The biguanides metformin and phenformin inhibit angiogenesis, local and metastatic growth of breast cancer by targeting both neoplastic and microenvironment cells. Int J Cancer 2015, 136(6):E534–544.

18. Zhu P, Davis M, Blackwelder AJ, Bachman N, Liu B, Edgerton S, Williams LL, Thor AD, Yang X: Metformin selectively targets tumor-initiating cells in ErbB2-overexpressing breast cancer models. Cancer Prev Res (Phila) 2014, 7(2):199–210.

19. Thompson MD, Grubbs CJ, Bode AM, Reid JM, McGovern R, Bernard PS, Stijleman IJ, Green JE, Bennett C, Juliana MM et al: Lack of effect of metformin on mammary carcinogenesis in nondiabetic rat and mouse models. Cancer Prev Res (Phila) 2015, 8(3):231–239.

20. Anisimov V, Egormin P, Bershtein L, Zabezhinskii M, Piskunova T, Popovich I, Semenchenko A: Metformin decelerates aging and development of mammary tumors in HER-2/neu transgenic mice. Bulletin of experimental biology and medicine 2005, 139(6):721–723.

21. Barbosa LB, Gama IRS, Vasconcelos NBR, Santos EAD, Ataide-Silva T, Ferreira HDS: Dietary patterns according to gender and ethnicity associated with metabolic syndrome: a systematic review and meta-analysis. Cien Saude Colet 2024, 29(10):e03662023.

22. Cejuela M, Martin-Castillo B, Menendez JA, Pernas S: Metformin and Breast Cancer: Where Are We Now? Int J Mol Sci 2022, 23(5).

23. Srinivasan M, Arzoun H, Gk LB, Thangaraj SR: A Systematic Review: Does Insulin Resistance Affect the Risk and Survival Outcome of Breast Cancer in Women? Cureus 2022, 14(1):e21712.

24. Peeters PJ, Bazelier MT, Vestergaard P, Leufkens HG, Schmidt MK, de Vries F, De Bruin ML: Use of metformin and survival of diabetic women with breast cancer. Curr Drug Saf 2013, 8(5):357–363.

25. Chang MC, Eslami Z, Ennis M, Goodwin PJ: Crown-like structures in breast adipose tissue of breast cancer patients: associations with CD68 expression, obesity, metabolic factors and prognosis. NPJ Breast Cancer 2021, 7(1):97.

