## Supplemental Tables for "Targeting the menopause transition with metformin improves breast cancer outcomes, but discontinuation has deleterious effects on metabolic health: Findings from a preclinical model of postmenopausal breast cancer"

**Supplemental Table 1**. Key characteristics of the rat model used in this study.

| Parameter | Details |
|---|---|
| Rodents | Female Wistar rats<br>(5 wks. of age, 87-148g; Charles River Laboratories, Wilmington, MA) |
| Caging | Individually housed in wire-bottomed cages to decrease physical activity |
| Environment | 22-24°C with a 12-h/12-h light-dark cycle |
| Diet | *Ad libitum* high-fat diet (46% kcal fat, Research Diets #12344) with free access to water |
| Tumor Initiation | Single intraperitoneal injection of 1-methyl-1-nitrosourea<br>(50 mg/kg; #MRI-340, MRI Global, Kansas City, MO) at 55 ± 2 days of age |

**Supplemental Table 2**. Histology antibodies and kits used to measure plasma metabolites in this study.

| Plasma Metabolite | Details |
|---|---|
| Insulin | ELISA Alpco 80-INSRT-E01, Salem, NH |
| Glucose | Colorimetric assay TR15421, Thermo Fisher Scientific, Waltham, MA |
| Triglycerides | Colorimetric assay TR22321, Thermo Fisher Scientific, Waltham, MA |
| Total cholesterol | Colorimetric assay TR13521, Thermo Fisher Scientific, Waltham, MA |
| Non-esterified fatty acids | Colorimetric assay, Wako Chemicals USA, Richmond, VA |

**Supplemental Table 3**. Body composition at study end, including total body mass, total lean mass, total fat mass, mass of individual adipose depots (mesenteric, retroperitoneal, and gonadal), and liver fat mass. Data are presented as mean ± standard error of the mean.

| Treatment | Control | $MET_{0\text{-}8wks}$ | $MET_{0\text{-}4wks}$ | $MET_{4\text{-}8wks}$ |
|---|---|---|---|---|
| Body Mass (g) | 396.2 ± 22.2 | 374.3 ± 21.9 | 402.0 ± 33.9 | 357.1 ± 20.7 |
| Fat Mass (g) | 137.6 ± 15.2 | 116.7 ± 13.2 | 148.1 ± 23.3 | 103.8 ± 9.9 |
| Lean Mass (g) | 228.9 ± 7.4 | 227.0 ± 10.2 | 224.2 ± 10.9 | 224.6 ± 11.7 |
| Mesenteric (g) | 6.0 ± 0.3 | 6.1 ± 0.3 | 6.0 ± 0.6 | 5.7 ± 0.4 |
| Retroperitoneal (g) | 11.7 ± 0.8 | 11.9 ± 0.6 | 12.5 ± 1.2 | 12.9 ± 1.1 |
| Gonadal (g) | 10.6 ± 0.6 | 11.3 ± 0.7 | 10.5 ± 0.7 | 10.6 ± 0.7 |
| Liver Fat (g) | 1.1 ± 0.1 | 1.2 ± 0.1 | 1.0 ± 0.2 | 1.0 ± 0.1 |

**Supplemental Table 4**. The concentration of plasma metabolites measured at study end. Data are presented as mean ± standard error of the mean.

| Treatment | Control | | | $MET_{0\text{-}8wks}$ | | | $MET_{0\text{-}4wks}$ | | | $MET_{4\text{-}8wks}$ | | |
|---|---|---|---|---|---|---|---|---|---|---|---|---|
| Glucose (mM) | 10.5 | ± | 0.6 | 10.9 | ± | 0.6 | 11.0 | ± | 0.4 | 10.3 | ± | 0.6 |
| NEFA (µM) | 697 | ± | 56 | 683 | ± | 43 | 710 | ± | 66 | 661 | ± | 55 |
| Cholesterol (mM) | 1.9 | ± | 0.2 | 1.8 | ± | 0.1 | 2.4 | ± | 0.4 | 1.6 | ± | 0.2 |
| Triglycerides (mM) | 0.5 | ± | 0 | 0.5 | ± | 0.1 | 0.5 | ± | 0.1 | 0.5 | ± | 0.1 |
| Leptin (pg/mL) | 6464 | ± | 1022 | 4395 | ± | 713 | 5935 | ± | 1252 | 2979 | ± | 536 |
